# Agonistic character displacement of genetically based male color patterns across darters

**DOI:** 10.1101/338715

**Authors:** Rachel L. Moran, Rebecca C. Fuller

## Abstract

A growing number of studies are demonstrating that interspecific male-male competitive interactions can promote trait evolution and contribute to speciation. Agonistic character displacement (ACD) occurs when selection to avoid maladaptive interspecific aggression leads to the evolution of agonistic signals and/or associated behavioral biases in sympatry. Here we test for a pattern consistent with ACD in male color pattern in darters (Percidae: *Etheostoma*). Despite the presence of traditional sex roles and sexual dimorphism, male color pattern has been shown to function in male-male competition in several darter species, rather than female mating preferences. A pattern consistent with divergent ACD in male behavioral biases has also been documented in darters. Males bias their aggression towards conspecific over heterospecific males in sympatry but not in allopatry. Here we use a common garden approach to show that differences in male color pattern among four closely related darter species are genetically based. We also demonstrate that male color pattern exhibits enhanced differences in sympatric compared to allopatric populations of two darter species. This study provides evidence that interspecific male-male aggressive interactions alone can promote elaborate male signal evolution both between and within species. We discuss the implications this has for male-driven ACD and cascade ACD.

## Introduction

Evolutionary biologists have long been interested in secondary contact events between previously allopatric lineages because they provide valuable insight into the process of speciation. Secondary contact can result in a variety of outcomes depending on the degree of reproductive isolation that has accrued [1–3]. For example, exploitative competition over shared resources might cause one lineage to go locally extinct. Another possibility is that the two lineages freely hybridize upon secondary contact and collapse into a hybrid swarm. Alternatively, selection against maladaptive hybridization between lineages can promote the evolution of reproductive character displacement (RCD), thereby finalizing the speciation process in sympatry. RCD occurs when selection to avoid interspecific mating results in the evolution of a mating traits (signals and/or preferences) [1,4]. Studies of RCD have focused largely on the evolution of female mating preferences and associated male traits (reviewed in 5). However, male mating preferences for female traits can also promote speciation via RCD [6–11]. Furthermore, a growing number of studies indicate that interspecific male-male competitive interactions can influence trait divergence and speciation in sympatry via agonistic character displacement [12–16]. Similar to RCD, ACD occurs when selection to avoid interspecific fighting results in the evolution of competitive traits (signals and/or aggression biases) [17,18]. Both RCD and ACD can result in a pattern of enhanced trait divergence between species in sympatry compared to allopatry.

When gene flow among populations within a species is low, RCD and ACD can incidentally cause mismatches among populations within a species in traits associated with mate/competitor evaluation [19,20]. The evolution of trait divergence among allopatric populations as a correlated effect of character displacement between sympatric species is termed “cascade” character displacement. Cascade RCD can cause increased behavioral isolation among populations within species. Cascade ACD can alter the likelihood of competitive interactions in secondary contact. Although cascade RCD has been demonstrated in a variety of taxa [21,22], darters (Percidae: *Etheostoma*) represent the only documented example of cascade ACD [16].

This study tests for divergent ACD in a male color pattern in darters, a diverse group of North American stream fishes. Verifying that the evolution of a given signal trait is a product of divergent ACD requires demonstrating: (1) that the signal functions in competitive interactions (rather than male-female mating interactions), (2) that the signal is genetically based and not due to environmental differences between sympatry and allopatry, and (3) that a geographic pattern of enhanced signal divergence between species in sympatry compared to allopatry is present [5,17,18]. Several recent studies have shown that male color pattern functions in male-male competition in darters. Within species, aspects of male color pattern predict a male’s ability to guard a female from rival males and consequently correlate with reproductive success [23]. Male color pattern also functions in male discrimination between conspecific versus heterospecific male competitors [24–26]. Furthermore, there is evidence for cascade RCD and cascade ACD because RCD and ACD between rainbow darters and the orangethroat clade leads to heightened isolation between allopatric orangethroat species (see Study System below). Here, we use a common garden approach to ask whether differences in male color pattern present among four closely related species of darters are genetically based. We then compare multivariate measurements of male color pattern in sympatric and allopatric population pairs in two darter species to test whether color pattern divergence is enhanced in sympatry compared to allopatry. This study provides important insight into the evolution of an elaborate sexually dimorphic color trait in a highly diverse group of vertebrates with traditional sex roles but no apparent female mating preferences. Our results demonstrate how interspecific male-male competition can lead to color pattern divergence between and within species and has implications for RCD, ACD, cascade RCD, and cascade ACD in darters.

## Methods

### Study System

This study focuses on two groups of darters: the orangethroat darter clade (*Etheostoma: Ceasia*) and the rainbow darter (*Etheostoma caeruleum*). The orangethroat darter clade includes 15 recently-diverged allopatric species. These new species have been diagnosed over the past several decades based largely on qualitative differences in male color pattern among populations in different drainages [27,28]. Recent research suggests that the dramatic diversification within the orangethroat darter clade may be driven by selection against reproductive and agonistic interactions with the rainbow darter. Thirteen out of the 15 species within the orangethroat clade occur in sympatry with the rainbow darter. These species hybridize at low levels in sympatry, and a substantial amount of postzygotic isolation is present in the form of male-skewed F1 hybrid sex ratios and high levels of backcross hybrid inviability [29]. When orangethroat and rainbow darters co-occur with one another, males exert strong preferences for mating with conspecific over heterospecific females and bias their aggression towards conspecific over heterospecific males [26]. Such preferences are absent in orangethroat and rainbow darters when they occur in allopatry with respect to one another [16]. Thus, it appears that selection to avoid costly interspecific interactions has led to male-driven RCD and ACD in sympatry between orangethroat and rainbow darters. Furthermore, orangethroat darter males show enhanced preferences for mating and fighting with conspecifics over individuals from other closely related species within the orangethroat clade only when they co-occur in sympatry with rainbow darters [16]. This suggests that RCD and ACD between orangethroat and rainbow darters has incidentally led to trait evolution and behavioral isolation among lineages within the orangethroat darter clade (i.e., cascade RCD and cascade ACD).

### Common Garden Study

Our goal here was to test whether color pattern differences present among species within the orangethroat clade are genetically based. We chose to focus on four species in the orangethroat clade that were recently shown to differ quantitatively from one another in the color pattern of wild-caught males: the orangethroat darter (*Etheostoma spectabile*), the strawberry darter (*E. fragi*), the current darter (*E. uniporum*), and the brook darter (*E. burri*) (Figure 1A) [26]. In March 2015, adult male and female fish from one population of each of the four species were collected using a kick seine (locations shown in Table S1). Fish were transported in aerated buckets back to the University of Illinois at Urbana-Champaign, sorted by sex and species, and maintained in 75.7 L stock tanks. For each species, we set up 37.9 L breeding tanks that contained a conspecific pair of one male and one female. We created three to four replicate crosses (i.e., families) for each of the four species. Breeding tanks were filled with three to five cm of naturally colored aquarium gravel. All stock and breeding tanks contained a sponge filter and tap water treated with dechlorinator. Tanks were maintained in the same room at 19° C under fluorescent lighting set to mimic the natural photoperiod. Fish were fed frozen bloodworms daily *ad libitum*.

**Figure 1.**
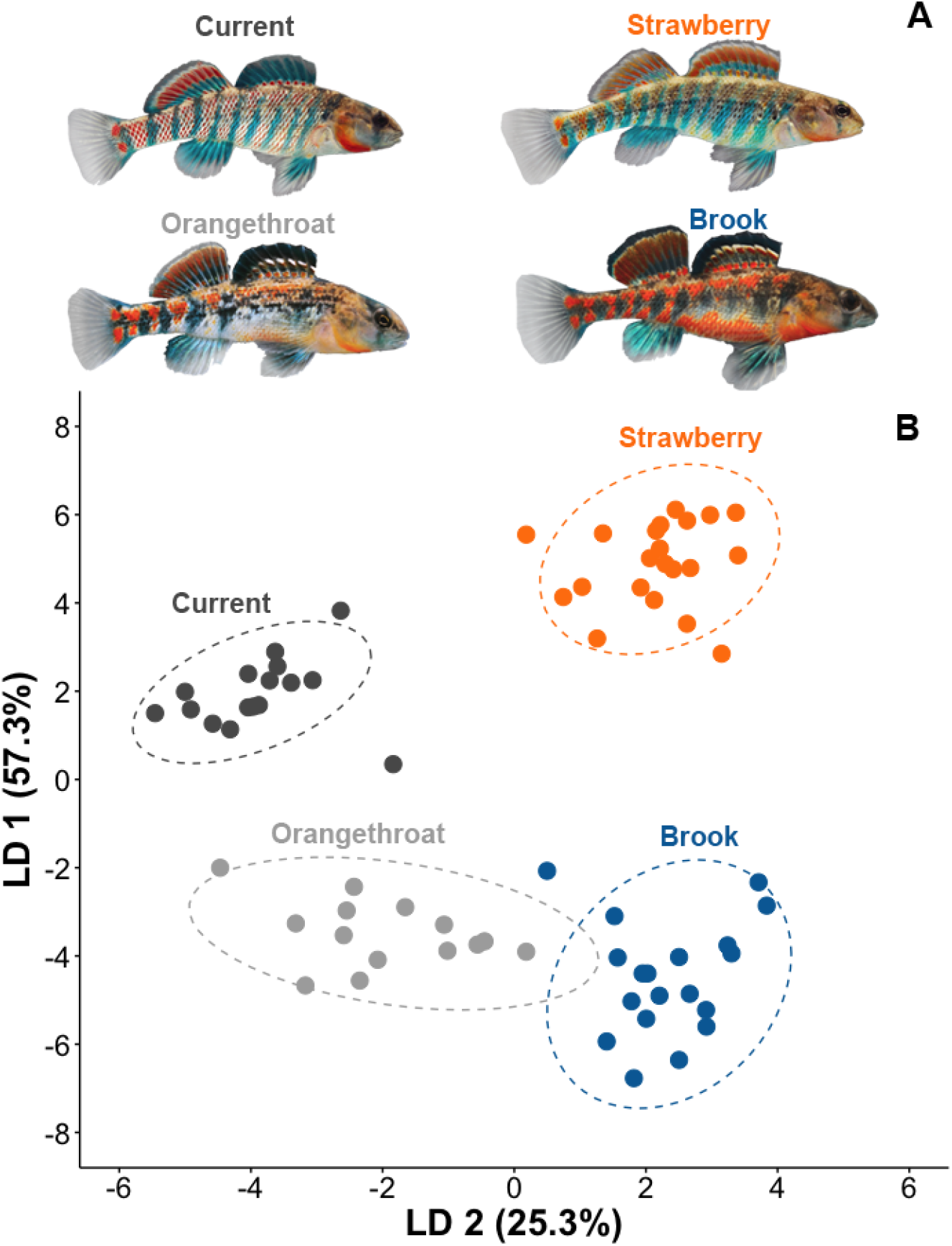
(A) Representative example of male color pattern in strawberry, current, brook, and orangethroat darters. (B) Biplot of the first two LDs obtained from the LDA on male color pattern in fish from the common garden study. Ellipses represent 95% CI.

Eggs were collected from breeding tanks using a gravel siphon every 1-3 days for a period of one month. All collected eggs were placed in 0.5 L plastic tubs filled with water treated with methylene blue to prevent fungal growth. Offspring from the same family were kept together. After hatching, fry were transferred to a 1 L plastic tub and fed live brine shrimp nauplii every other day. At approximately 1 month of age, fry were large enough for frozen daphnia to be incorporated into their diet. At approximately three months of age, we transitioned to feeding the fry daily with frozen bloodworms. At this time, all families were transferred to 2.5 L tanks. At one year of age, fish were transferred to 37.9 L tanks, and at two years of age they were transferred to 75.7 L tanks. Offspring from all families were housed in the same room at 19° C under fluorescent lighting that mimicked the natural photoperiod.

At approximately three years of age, the lab-raised offspring from each of the four species had reached adult size and males had developed adult breeding coloration. At this time, males from each family were photographed with a Nikon Coolpix D3300 digital camera (mean ± SE males per family = 5.5 ± 0.6). Photographs were taken under fluorescent lighting with the camera’s factory setting for fluorescent light. Prior to photographing, fish were lightly anesthetized using 0.03g/L of MS-222 and were then placed in a petri dish filled with treated water. An X-rite ColorChecker Mini Chart (Grand Rapids, MI) was in each photograph for color correction and standardization with the inCamera 4.5 plug-in for Adobe Photoshop CC (Adobe Systems Inc., San Jose, CA). We also included a ruler in each photograph, which we used to measure the standard length of each fish (i.e., tip of snout to end of caudle peduncle) to the nearest mm in ImageJ (version 1.50c4) [30].

Males from all species within the orangethroat clade exhibit red and blue banding on the lateral side of the body and on the two dorsal fins (Figure 1A). To quantify any differences in male color pattern that were present at the species level, we focused our analyses on aspects of male color pattern that have been shown previously to contribute to variation among these species [26]. We measured RGB values for both the red and blue coloration on the body, as well as the proportion of red and blue coloration present on the body and fins. The dropper tool in Adobe Photoshop was used to measure RGB values, which vary 0-255 for each of the three color channels (i.e., red, green, and blue). An RGB value of 0,0,0 represents black and 255,255,255 represents white. We recorded the three values associated with RGB in both the red and blue portion of the color pattern (resulting in six RGB variables total) on the posterior half of the lateral side of each fish, near the caudal peduncle. The dropper tool was set to sample a 3×3 pixel area within a given color patch. Each location was measured three times, and the average of these measurements was used for each fish in the multivariate analysis. We used ImageJ to measure the proportion of red and blue on the body and fins as described in Moran et al. [26,29]. Briefly, the perimeter of each fin and the body were traced separately using the polygon selection tool and the areas for each part of the fish were calculated with the histogram function. We then isolated the red and blue pixels using the Threshold Colour Plugin with the color channel set to CIE *Lab*. Once the red or blue pixels were isolated, we made the image binary and counted the number of black pixels in the regions corresponding to the fins and the body. We measured the proportion of red and the proportion of blue present on the lateral side of the body and the two dorsal fins, for a total of six color proportion variables per fish.

All statistical analyses were performed in R (version 3.4.4). We first conducted a two-factor nested MANOVA to examine whether color pattern differed significantly among species and among families (i.e., replicate crosses within a species). Each of the 12 color pattern variables served as dependent variables in this analysis, with species and family (nested within species) included as factors. We also conducted two-factor nested ANOVAs for each dependent variable to determine whether significant differences existed among families and species. Because size (i.e., standard length in mm) varied among families (F_12,58_ = 11.72, P < 0.00001), it was included as covariate in the MANOVA and ANOVAs. However, preliminary analyses indicated no effect of size on color pattern differences among individuals. We therefore excluded size from subsequent analyses. We next used Linear Discriminate Analysis (LDA) to reduce the dimensionality of the color data set and to identify which color variables contribute most to differences among species. We used the lda function of the MASS package [31]. The 12 color pattern measurements served as dependent variables, and species served as the categorical predictor variable. We then used the Anova function of the car package [32] to conduct nested two-factor ANOVAs with species and family (nested within species) as factors and individual linear discriminate scores as the dependent variable. We conducted separate ANOVAs for both of the first two linear discriminants. Post-hoc pairwise comparisons were conducted among species using Tukey’s tests with the glht function in the multcomp package [33].

### ACD Study

Here our goal was to quantify male color pattern variation in wild-caught sympatric and allopatric populations of the orangethroat darter and the rainbow darter to test for a pattern consistent with divergent ACD. The orangethroat darter is the only species within the orangethroat clade to occur both in sympatry and in allopatry with respect to the rainbow darter. Previous studies have shown that aspects of male color pattern differ quantitatively between sympatric orangethroat and rainbow darters [26,29], and that color pattern is variable across populations within species [34]. Divergent ACD in male color pattern predicts: (1) enhanced differentiation between species in sympatry compared to allopatry, and (2) differentiation between sympatric and allopatric populations within species.

Adult orangethroat and rainbow darter males were collected with a kick seine in March 2016 from one sympatric and one allopatric population of each species (for a total of four “groups”) (Figure 2A, Table S2). We took digital photographs of 10 males from each group, for a total of 40 fish, as described above for the common garden study. Size did not vary among the four groups (ANOVA: F_3,36_ = 1.88, P = 0.15).

**Figure 2.**
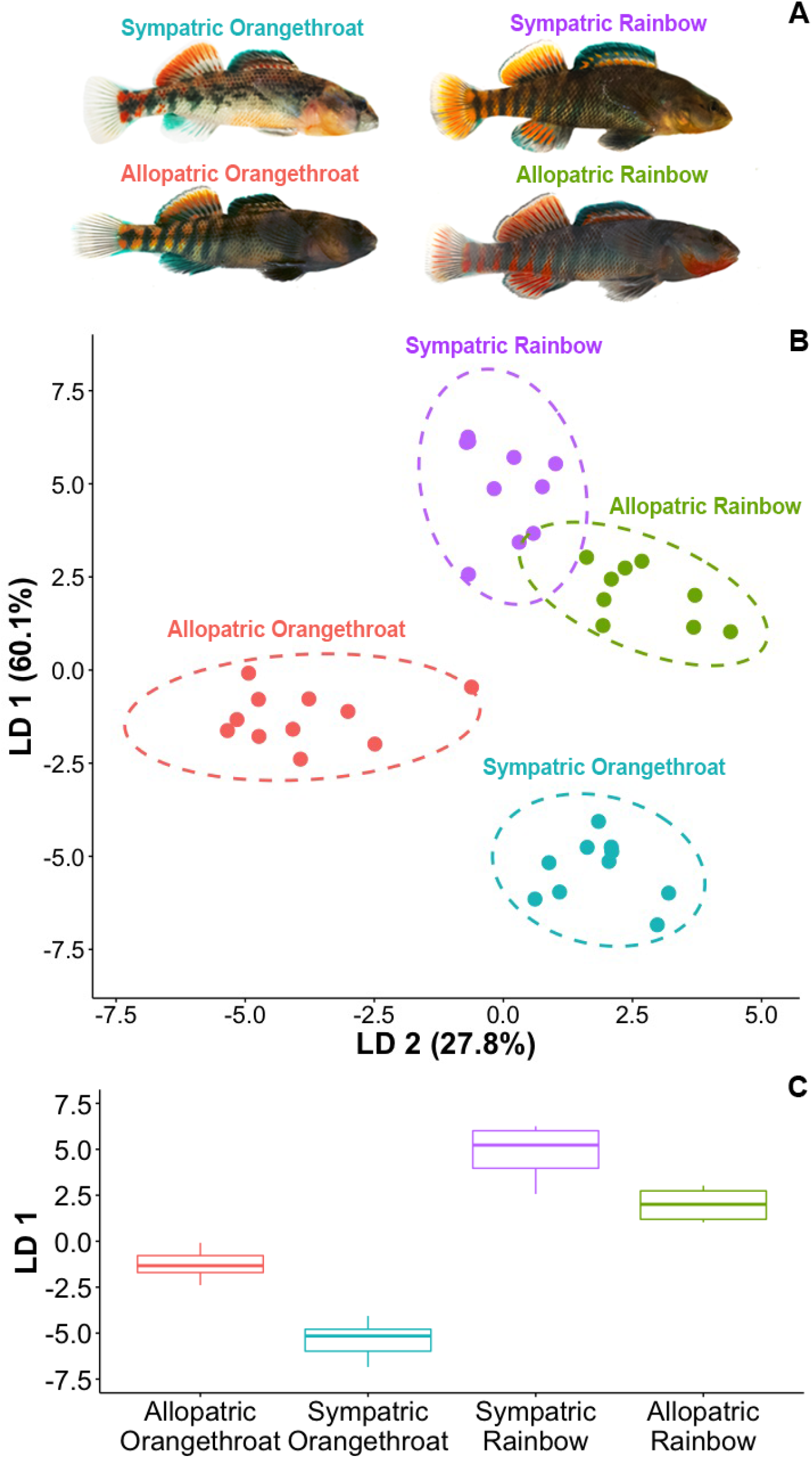
(A) Representative example of male color pattern in sympatric orangethroat, sympatric rainbow, allopatric orangethroat, and allopatric rainbow darters. (B) Biplot of the first two LDs obtained from the LDA on male color pattern in fish from the ACD study. Ellipses represent 95% CI. (C) Boxplots of LD2 scores from the LDA on male color pattern in fish from the ACD study.

Orangethroat and rainbow darters are both characterized by a male nuptial color pattern that consists of red and blue banding on the lateral sides and dorsal fins. Despite their superficial similarities, their color patterns differ in a few consistent ways (Figure 2A). Orangethroat darters lack red coloration on their anal fins, caudal fins, and pectoral fins, but rainbow darters do not. There are also apparent differences in the amount of red and blue banding across the lateral portion of the fish. To quantify variation in male color pattern between and within species, we followed the methods described above for the common garden study. In addition, we measured the proportion of red coloration present on the caudal fin and the proportion of red and proportion of blue coloration present on the anal fin for each fish.

All analyses were conducted in R using the packages described above. We first conducted a two-factor nested MANOVA to examine whether species and geography (i.e., sympatric or allopatric) contributed to differences in male color pattern among groups. Each of the 16 color pattern variables served as dependent variables in this analysis, with species and geography (nested within species) included as factors. We also conducted two-factor nested ANOVAs for each dependent variable with species and geography as factors. We then used LDA to facilitate pairwise comparisons and to identify which variables contribute most to differences among groups. Here, group (i.e., sympatric rainbow, allopatric rainbow, sympatric orangethroat, or allopatric orangethroat) served as the categorical predictor variable and the color measurements served as dependent variables. Finally, to ask whether individuals’ scores for the first two LDs differed among groups, we used nested two-factor ANOVAs. We included the score for the first and second LDs as the dependent variable (in two separate analyses, one for each LD). Species and geography (nested within species) were included as factors. We made post-hoc pairwise comparisons among groups using Tukey’s tests.

## Results

### Common Garden Study

The MANOVA revealed that species identity and family (i.e., replicate cross within a species) both significantly contributed to differences in male color pattern among lab-raised fish (Table 1). There was no effect of size (standard length in mm) on differences in color pattern among individuals (Table 1). ANOVAs indicated that the values for nearly every variable differed significantly as a function of species identity or due to an interaction between family and species (Table S3). The one exception was the proportion of blue present on the second dorsal fin, which varied among families within species but not among species.

**Table 1.** Results of two-factor nested MANOVA on male color pattern in fish from the common garden study. Species and family (nested within species) were included as factors and size (standard length in mm) was included as a covariate.

| Variable | Df | Pillai | approx F | num Df | den Df | P |
| --- | --- | --- | --- | --- | --- | --- |
| Species | 1, 62 | 0.61 | 6.54 | 12 | 51 | <0.00001 |
| Family | 3, 62 | 2.60 | 28.86 | 36 | 159 | <0.00001 |
| Size | 1, 62 | 0.31 | 1.92 | 12 | 51 | 0.054 |
| Family*Species | 3, 62 | 1.10 | 2.54 | 36 | 159 | <0.00001 |

The LDA reduced the dimensionality of the color pattern measurements into three LDs, with the first two LDs explaining 86.6% cumulative variation among groups (LD1: 57.3%, LD2: 25.3%, LD3: 17.4%). Figure 1B shows a biplot comparing the scores for LD1 versus LD2 for each individual, grouped by species. The color pattern proportion measurements had higher loadings (i.e., associations) with all three LDs compared to the RGB data, suggesting that differences in the proportion of red and blue coloration on the body and fins is a good predictor of species.

The proportion of red on the first and second dorsal fins and the proportion of blue on the first dorsal fin had the highest loadings for LD1. LD2 was associated with the proportion of red on both dorsal fins in addition to the proportion of red present on the body.

ANOVAs on LD1 and LD2 revealed significant effects of species (LD1: F_3,63_ = 35.70, P < 0.00001; LD2: F_3,63_ = 15.60, P < 0.00001) but not family (nested within species) (LD1: F_1,63_ = 2.90, P = 0.09; LD2: F_1,63_ = 1.14, P = 0.29). There was no interaction between species and family for either analysis (LD1: F_3,63_ = 0.73, P = 0.54; LD2: F_3,63_ = 0.53, P = 0.66). Post-hoc Tukey’s tests indicated that all species differed significantly from one another in scores for LD1 and/or LD2 (i.e., no pair of species overlapped in scores for both LD1 and LD2) (Tables S4).

### ACD Study

Our MANOVA on variation in male color pattern among groups (i.e., allopatric rainbow, allopatric orangethroat, sympatric rainbow, and sympatric orangethroat) revealed an interaction between species identity (orangethroat or rainbow) and geography (sympatric or allopatric, nested within species), but not size (Table 2). ANOVAs indicated that the values for nearly every variable differed significantly between sympatric and allopatric populations within species geography or due to an interaction between geography and species (Table S5). The red (R) value for the blue coloration, the proportion of blue coloration on the body, and the proportion of red coloration on the body and anal fin varied between species but was not associated with geography.

**Table 2.**
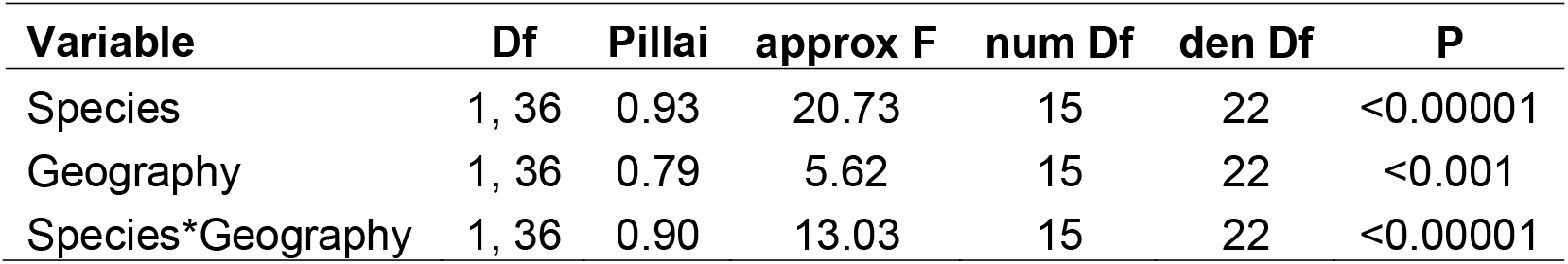
Results of two-factor nested MANOVA on male color pattern in fish from ACD study. Species and geography (nested within species) were included as factors.

LDA identified three LDs that predicted differences among groups, with the first two LDs explaining 87.9% cumulative variation among groups (LD1: 60.1%; LD2: 27.8%; LD3: 12.1%). Figure 2B shows a biplot comparing the scores for LD1 versus LD2 for each individual. The color pattern proportion measurements had higher loadings (i.e., associations) with all three LDs compared to the RGB data, suggesting that differences in the proportion of red and blue coloration on the body and fins is a good predictor of species and geographic relationship between groups.

Contrasting patterns were present in scores for the first two LDs across groups. A pattern consistent with divergent character displacement was evident from LD1 (Figure 2B,C). Scores for LD1 showed a closer association between allopatric fish compared to sympatric fish of both species. This pattern was mainly driven by differences between sympatric and allopatric populations of orangethroat darters. LD1 was most strongly associated with the proportion of red coloration present on the anal fin, caudal fin, and body. Conversely, sympatric males of both species were grouped more closely along LD2 compared to allopatric males of both species (Figures 2B, S1). LD2 was most closely associated with the proportion of blue coloration on the first and second dorsal fin. This suggest that traits corresponding with LD2 may be associated with sharing a common environment and/or gene flow.

ANOVAs for both LD1 and LD2 indicated an interaction between species and geography (nested within species) (LD1: F_1,36_ = 127.78, P < 0.0001; LD2: F_1,36_ = 178.55, P < 0.0001). Post-hoc pairwise comparisons with Tukey’s tests revealed significant differences among all groups in scores for LD1 (Table S6A). Only one pairing did not differ significantly from one another in scores for LD2: allopatric rainbow darters and sympatric orangethroat darters (Table S6B).

## Discussion

A growing body of literature suggests that interspecific reproductive and aggressive interactions play a surprisingly large role in speciation [17,18]. Interspecific interactions can have broad implications for speciation by directly promoting enhanced behavioral isolation in sympatry and indirectly promoting the evolution of trait divergence and behavioral isolation among allopatric lineages [21,22]. In this study we demonstrated that color pattern differences present in nature among recently diverged allopatric lineages within the orangethroat clade are genetically based. Additionally, we observed a pattern of enhanced divergence in male agonistic coloration in sympatry (compared to allopatry) between populations of the orangethroat darter and the more distantly related rainbow darter, consistent with divergent ACD. These results have significant implications for our understanding of speciation and diversification in one of the most diverse groups of vertebrates in North America. More generally, this study provides important insight into the evolution of ACD and cascade ACD in male agonistic signals and response to those signals.

A unique aspect of this study system is that evolution of elaborate male nuptial coloration appears to be driven entirely by male-male interactions between and within species, despite the presence of traditional sex roles. Previous studies on orangethroat and rainbow darters have demonstrated that male coloration functions in male-male competition over access to females within species [23], and that male aggressive response towards heterospecific males increases with increasing color pattern similarity between species [26]. Conversely, studies have consistently failed to detect female preferences associated with variable aspects of male color pattern within or between species [16, 24, 26,35, 36]. Here we demonstrated that some male color traits (i.e., those associated with LD1 in the ACD study: proportion of red coloration on the anal fin, caudal fin, and body) show a clear pattern of divergent character displacement between sympatric orangethroat and rainbow darter populations (Figure 2C). Thus, the results of the present study together with previous behavioral experiments suggest that male color pattern evolution in sympatry is driven by divergent ACD between orangethroat and rainbow darters.

We also found that some aspects of male color pattern (i.e., those associated with LD2 in the ACD study: proportion of blue coloration on the first and second dorsal fin) appear to be more strongly correlated with a common environment and/or gene flow, and do not show a pattern consist with divergent ACD between orangethroat and rainbow darters (Figures 2B,S1). Theoretically, the greater similarity in sympatry compared to allopatry in some color traits may be due to three different phenomena: introgression due to hybridization, local adaptation to a common environment, or phenotypic plasticity due to sharing a common environment. We doubt that phenotypic plasticity accounts for the convergence in color proportion traits on the dorsal fins. Clearly, there are some types of color traits that are plastic. Red coloration in darters is carotenoid based [34], which suggests it may be linked to diet [37,38]. In rainbow darters, spectral properties of red coloration are associated with parasite load [39]. Additionally, blue and black coloration present on the side of the body and head can vary rapidly in these species when males escalate aggression (R. Moran pers. obs.). However, these phenomena should affect the red and blue hues and their associated RGB values. The present study has demonstrated that variation in RBG values account for little of the total variation present between sympatric and allopatric populations/species. Instead, the proportion of red and blue coloration present on the body and fins strongly predict both species identity and geographic relationship between species. The results of our common garden study in combination with another recent study examining male color pattern in orangethroat darters, rainbow darters, and their hybrids [29] provide strong evidence that these color elements are largely genetic in nature.

The other two phenomena that can potentially account for the convergence in some color traits are hybridization and local adaptation. Of these two possibilities, we suspect that hybridization is more likely for two reasons. First, hybridization is ongoing in at least three different contact zones. Moran et al. [26,29] and Bossu and Near [40] have shown that F1 hybrids between rainbow darters and three different orangethroat clade species are present in natural populations. In addition, the traits that are most strongly implicated in species-specific differences between orangethroat and rainbow darters (i.e., the proportion of red and blue coloration on the body, anal fin, and caudal fin) have intermediate values in F1 hybrid males [29], suggesting that introgression can cause increased trait similarity between species. Second, although large-scale transitions between genera and sub-genera are associated with ecological divergence in darters [41,42], there is strong evidence that differences in male color pattern among more closely related species are primarily driven by intrasexual selection rather than ecological differences [24,26,43,44].

Importantly, the findings of this study drastically change how we think about the evolution of male color pattern and speciation in darters. Sexual selection in the form of female mating preference for male color traits was long thought to be the primary catalyst of speciation in these fish. We have demonstrated that divergence in male color pattern both between species and among populations within species is promoted in sympatry between congeners. This observation reflects our previously documented behavioral patterns of ACD between orangethroat and rainbow darters and cascade ACD among species in the orangethroat clade. Divergence in male color traits between closely related species within the orangethroat clade that occur in sympatry with rainbow darters (and thus undergo ACD) has resulted in enhanced male competitor bias between species [16,26]. This is consistent with cascade ACD in both male agonistic signals and behavioral response to those signals (sensu “convergent sympatry effects” of character displacement) [20]. It remains to be tested whether the divergence in male color pattern traits that we observed between populations within the orangethroat darter (and/or within the rainbow darter) also confer behavioral biases among populations within species (which would indicate “sympatry-allopatry effects” of character displacement) [20].

Our findings also have implications for the evolution of behavioral isolation via RCD and cascade RCD in this system. Our current hypothesis is that strong selection to avoid maladaptive hybridization after secondary contact (i.e., reinforcement) leads to RCD in male mating preferences and strong behavioral isolation between species [29]. As a result, females of both species are not a shared resource among males of both species in sympatry, which could cause interspecific male-male aggression over females to be maladaptive. This should promote ACD in male aggressive biases (to avoid needless interspecific aggression), allowing these species to co-occur in close proximity on the breeding grounds and in turn increasing the potential for hybridization. In this manner RCD and ACD may act in a positive feedback loop, mutually strengthening divergence in both mating and fighting traits in males.

To conclude, the results of the present study demonstrate that interspecific interactions in sympatry may play a larger role than previously thought in promoting the diversification of male secondary sex traits both between and within species. Evidence is now growing that female mating preferences are absent or lower compared to male mate preferences in many species of darters. Instead, it appears that male mating preferences and aggressive biases drive trait evolution between and within species, despite the presence of elaborate male secondary sex traits and traditional sex roles.

## Author Contributions

RLM drafted the manuscript, carried out the studies, and analyzed the data. RCF made revisions to the manuscript and assisted RLM with design of the study. Both authors gave final approval for the publication.

## Acknowledgements

The treatment of animals used in this study was in compliance with the University of Illinois Institutional Animal Care and Use Committee (IACUC) under protocols #14097 and #17031. Collection of wild fish was approved by the Illinois Department of Natural Resources under Scientific Collecting Permits A15.4035 and A16.4035, the Arkansas Game and Fish Commission under Scientific Collection Permit #032020141, the Missouri Department of Conservation under Wildlife Collector’s Permit #16392, and the Michigan Department of Natural Resources under a Scientific Collector’s Permit issued to RLM and RCF. We thank Lance Merry for photos of darters that appear in Figure 1A.

## References

1. Coyne J, Orr H. 2004 Speciation. Sunderland, MA.

2. Harrison RG. 1993 Hybrid Zones and the Evolutionary Process. Oxford University Press, Oxford.

3. Liou LW, Price TD. 1994 Speciation by Reinforcement of Premating Isolation. Evolution. 48, 1451. (doi:10.2307/2410239)

4. Brown WL, Wilson EO. 1956 Character Displacement. Syst. Zool. 5, 49. (doi:10.2307/2411924)

5. Pfennig D, Pfennig K. 2012 Evolution’s wedge: competition and the origins of diversity. University of California Press, Berkeley, CA.

6. Servedio MR. 2007 Male versus female mate choice: Sexual selection and the evolution of species recognition via reinforcement. Evolution. 61, 2772–2789. (doi:10.1111/j.1558-5646.2007.00247.x)

7. Gabor CR, Ryan MJ. 2001 Geographical variation in reproductive character displacement in mate choice by male sailfin mollies. Proc. R. Soc. B Biol. Sci. 268, 1063–1070. (doi:10.1098/rspb.2001.1626)

8. Albert AAYK, Schluter D. 2004 Reproductive character displacement of male stickleback mate preference: reinforcement or direct selection? Evolution. 58, 1099–1107. (doi:10.1554/03-472)

9. Shine R, Phillips B, Waye H, Lemaster M, Mason RT. 2004 Species-isolating mechanisms in a mating system with male mate choice (garter snakes, Thamnophis spp.). Can. J. Zool. 82, 1091–1098. (doi:10.1139/z04-086)

10. Kozak GM, Roland G, Rankhorn C, Falater A, Berdan EL, Fuller RC. 2015 Behavioral Isolation due to Cascade Reinforcement in *Lucania* Killifish. Am. Nat. 185, 491–506. (doi:10.1086/680023)

11. Wiernasz DC. 1995 Male choice on the basis of female melanin pattern in Pieris butterflies. Anim. Behav. 49, 45–51. (doi:10.1016/0003-3472(95)80152-9)

12. Qvarnström A, Vallin N, Rudh A. 2012 The role of male contest competition over mates in speciation. Curr. Zool. 58, 493–509. (doi:10.1093/czoolo/58.3.493)

13. Vallin N, Rice AM, Bailey RI, Husby A, Qvarnström A. 2012 Positive feedback between ecological and reproductive character displacement in a young avian hybrid zone. Evolution. 66, 1167–1179. (doi:10.1111/j.1558-5646.2011.01518.x)

14. Drury JP, Grether GF. 2014 Interspecific aggression, not interspecific mating, drives character displacement in the wing coloration of male rubyspot damselflies (Hetaerina). Proc. R. Soc. B Biol. Sci. 281, 20141737. (doi:10.1098/rspb.2014.1737)

15. Okamoto KW, Grether GF. 2013 The evolution of species recognition in competitive and mating contexts: The relative efficacy of alternative mechanisms of character displacement. Ecol. Lett. 16, 670–678. (doi:10.1111/ele.12100)

16. Moran RL, Fuller RC. 2018 Male-driven reproductive and agonistic character displacement in darters and its implications for speciation in allopatry. Curr. Zool. 64, 101–113. (doi:10.1093/cz/zox069)

17. Grether GF, Losin N, Anderson CN, Okamoto K. 2009 The role of interspecific interference competition in character displacement and the evolution of competitor recognition. Biol. Rev. 84, 617–635. (doi:10.1111/j.1469-185X.2009.00089.x)

18. Grether GF, Peiman KS, Tobias JA, Robinson BW. 2017 Causes and consequences of behavioral interference between species. Trends Ecol. Evol. 32, 760–772. (doi:10.1016/j.tree.2017.07.004)

19. Yukilevich R, Aoki F. 2016 Is cascade reinforcement likely when sympatric and allopatric populations exchange migrants? Curr. Zool.

20. Comeault AA, Matute DR. 2016 Reinforcement’s incidental effects on reproductive isolation between conspecifics. Curr. Zool. 62, 135–143. (doi:10.1093/cz/zow002)

21. Ortiz-Barrientos D, Grealy A, Nosil P. 2009 The genetics and ecology of reinforcement: implications for the evolution of prezygotic isolation in sympatry and beyond. Ann. N. Y. Acad. Sci. 1168, 156–82. (doi:10.1111/j.1749-6632.2009.04919.x)

22. Hoskin CJ, Higgie M. 2010 Speciation via species interactions: The divergence of mating traits within species. Ecol. Lett. 13, 409–420. (doi:10.1111/j.1461-0248.2010.01448.x)

23. Zhou M, Fuller RC. 2016 Intrasexual competition underlies sexual selection on male breeding coloration in the orangethroat darter, *Etheostoma spectabile*. Ecol. Evol. 6, 3513–3522. (doi:10.1002/ece3.2136)

24. Zhou M, Loew ER, Fuller RC. 2015 Sexually asymmetric colour-based species discrimination in orangethroat darters. Anim. Behav. 106, 171–179. (doi:10.1016/j.anbehav.2015.05.016)

25. Martin MD, Mendelson TC. 2016 Male behaviour predicts trait divergence and the evolution of reproductive isolation in darters (Percidae: *Etheostoma*). Anim. Behav. 112, 179–186. (doi:10.1016/j.anbehav.2015.11.027)

26. Moran RL, Zhou M, Catchen JM, Fuller RC. 2017 Male and female contributions to behavioral isolation in darters as a function of genetic distance and color distance. Evolution. 71, 2428–2444. (doi:10.1111/evo.13321)

27. Ceas PA, Page LM. 1997 Systematic studies of the *Etheostoma spectabile* complex (Percidae; Subgenus *Oligocephalus*), with descriptions of four new species. Copeia, 496–522. (doi:10.2307/1447555)

28. Distler DA. 1968 Distribution and variation of *Etheostoma spectabile* (Agassiz) (Percidae, Teleostei). Univ. Kansas Sci. Bull. 48, 143–208.

29. Moran RL, Zhou M, Catchen JM, Fuller RC. 2018 Hybridization and postzygotic isolation promote reinforcement of male mating preferences in a diverse group of fishes with traditional sex roles. bioRxiv (doi:10.1101/325498)

30. Rasband W. 2011 ImageJ U. S. National Institutes of Health, Bethesda, Maryland, USA. https://ci.nii.ac.jp/naid/10030139275/ (accessed on 23 May 2018).

31. Ripley B, Venables B, Bates D, Hornik K, Gebhardt A. 2017 Package ‘MASS’.

32. Fox J. 2007 Package ‘car’. R Foundation for Statistical Computing.

33. Hothorn T, Bretz F, Westfall P, Heiberger R. 2017 Package ‘multcomp’.

34. Zhou M, Johnson AM, Fuller RC. 2014 Patterns of male breeding color variation differ across species, populations, and body size in rainbow and orangethroat darters. Copeia 2014, 297–308. (doi:10.1643/CI-12-103)

35. Pyron M. 1995 Mating patterns and a test for female mate choice in *Etheostoma spectabile* (Pisces, Percidae). Behav. Ecol. Sociobiol. 36, 407–412. (doi:10.1007/BF00177336)

36. Fuller RC. 2003 Disentangling female mate choice and male competition in the rainbow darter, Etheostoma caeruleum. Copeia 2003, 138–148.

37. Hill G, McGraw K. 2006 Bird coloration: mechanisms and measurements. Harvard University Press, Cambridge, MA.

38. Kodric-Brown A. 1989 Dietary carotenoids and male mating success in the guppy: an environmental component to female choice. Behav. Ecol. Sociobiol. 25, 393–401. (doi:10.1007/BF00300185)

39. Ciccotto PJ, Dresser DJ, Mendelson TC. 2014 Association between parasite load and orange, but not blue, male nuptial colouration in Etheostoma caeruleum. J. Fish Biol. 84, 1590–1598. (doi:10.1111/jfb.12361)

40. Bossu CM, Near TJ. 2013 Characterization of a contemporaneous hybrid zone between two darter species (*Etheostoma bison* and *E. caeruleum*) in the Buffalo River System. Genetica 141, 75–88. (doi:10.1007/s10709-013-9707-8)

41. Bossu CM, Near TJ. 2015 Ecological constraint and the evolution of sexual dichromatism in darters. Evolution. 69, 1219–1231. (doi:10.1111/evo.12655)

42. Ciccotto PJ, Mendelson TC. 2016 The ecological drivers of nuptial color evolution in darters (Percidae: Etheostomatinae). Evolution. 70, 745–756. (doi:10.1111/evo.12901)

43. Martin MD, Mendelson TC. 2014 Changes in sexual signals are greater than changes in ecological traits in a dichromatic group of fishes. Evolution. 68, 3618–3628. (doi:10.1111/evo.12509)

44. Martin MD, Mendelson TC. 2016 The accumulation of reproductive isolation in early stages of divergence supports a role for sexual selection. J. Evol. Biol. 29, 676–689. (doi:10.1111/jeb.12819)

